# Tumor extracellular vesicle RNA profiling predicts treatment response in pediatric diffuse midline glioma

**DOI:** 10.64898/2026.06.02.729542

**Authors:** Changheon Kim, Junbeom Kim, Sunjong Ji, Junhee Choi, Kiwan Bong, Ashley Pearson, Benison Lau, Carl J. Koschmann, Jouha Min

## Abstract

Detection of reliable markers of therapy response and drug resistance remains a major unmet need in brain cancer, as serial tumor biopsy is often not feasible. This challenge is particularly acute in diffuse midline glioma (DMG), a fatal pediatric brain tumor for which new targeted therapies are entering clinical use, yet tools for real-time molecular analysis of tumor evolution during therapy remain lacking. Here, we demonstrate that plasma tumor–derived extracellular vesicle (EV) profiling provides a minimally invasive and complementary approach for diagnosis and longitudinal molecular monitoring in H3K27M-mutant DMG. Across patient-derived tumor models and clinical plasma samples, EV mRNA levels correlated strongly with parental tumor transcriptomes. EV H3K27M mRNA enabled discrimination of DMG from non-DMG controls, and exploratory EV response-associated mRNAs were linked to radiographic response and progression-free survival in patients receiving ONC201. To enable tumor-selective EV enrichment and multiplexed molecular profiling within a clinically practical workflow, we developed a new integrated platform supporting same-day EV analysis from small plasma volumes. This work represents the first proof-of-concept demonstration of the diagnostic and treatment response relevance of tumor-derived EV RNA in pediatric DMG and establishes a generalizable framework for minimally invasive longitudinal molecular monitoring in diseases where tissue access is inherently limited.

## INTRODUCTION

Diffuse midline glioma (DMG), as defined in the 2021 World Health Organization (WHO) classification, is a highly aggressive grade IV glioma that encompasses most diffuse intrinsic pontine gliomas (DIPGs).^1, 2^ DMG predominantly affects children and young adults and has an exceptionally poor prognosis due to infiltrative growth in critical midline structures, which limits tissue access and precludes surgical resection.^3^ Most DMGs (∼80%) harbor a lysine-to-methionine substitution at position 27 of histone H3 (H3K27M), affecting the H3.1 or H3.3 variants.^4^ Patients with H3K27M-mutant tumors have a median survival of ∼9–12 months and a 5-year survival rate <10%.^5^ Radiation is the current standard of care but provides only transient benefits, while chemotherapy has not significantly improved survival.^6^

Recently, orally administered, blood-brain barrier penetrant imipridone compounds—including ONC201^7-9^ and its next-generation analog ONC206^10^—have emerged as promising targeted therapies for H3K27M-DMG. These agents act through antagonism of dopamine receptor D2 (DRD2) and activation of the mitochondrial protease ClpP, inducing cellular stress responses and tumor cell death.^11-14^ Clinical studies have reported impressive activity of ONC201 in DMG patients, including sustained (>2-year) radiographic responses in some cases,^15, 16^ leading to recent FDA approval as the first targeted therapy for this disease.^17^ However, therapeutic responses remain heterogeneous, and resistance frequently develops during treatment.

A major challenge in DMG management is the absence of minimally invasive and serially repeatable methods for real-time monitoring of evolving tumor state. Repeat biopsy is rarely feasible due to anatomical risk. MRI remains the clinical standard for response assessment but often lags behind molecular changes and may be confounded by pseudo-progression. Liquid biopsy approaches, including circulating tumor DNA (ctDNA)^18, 19^ and microRNA^20^ profiling, have shown feasibility in plasma and cerebrospinal fluid (CSF). However, ctDNA levels are often low in primary brain tumors, repeated CSF sampling is procedurally burdensome, and fragmented DNA provides limited insight into active transcriptional programs associated with therapeutic response and resistance.

Extracellular vesicles (EVs) represent a complementary and potentially useful biomarker modality for DMG. EVs are membrane-bound nanovesicles actively secreted by living cells— most notably by rapidly dividing tumor cells.^21, 22^ They are abundant and stable in bodily fluids, readily cross the blood-brain barrier, and carry diverse molecular cargo (e.g., proteins, nucleic acids, lipids) that reflects the molecular state of their parental cells.^23-28^ In contrast to fragmented genomic DNA, EV RNA may provide a dynamic readout of tumor biology during therapy. Although EVs have been investigated in adult gliomas and other malignancies,^29-34^ their use for plasma-based diagnosis and treatment-response monitoring in pediatric H3K27M-DMG remains largely unexplored.

Here, we demonstrate that circulating tumor-derived EV mRNA carries clinically relevant information in pediatric H3K27M-DMG. We show that EV H3K27M mRNA enables diagnostic discrimination of DMG from non-DMG healthy controls, and that exploratory EV response-associated mRNA levels (EGFR and FOXG1) are linked to progression-free survival (PFS) and radiographic response in patients receiving ONC201. To enable tumor-selective enrichment and multiplex RNA analysis from limited plasma volumes, we developed ExPLEX (Extracellular Vesicle Multi-PLEXer), a same-day assay workflow integrating immuno-enrichment, geometric multiplex encoding, and automated deep-learning-based decoding. Together, these findings extend EV RNA profiling to plasma-based molecular monitoring of pediatric DMG and support its use as a complementary strategy for longitudinal tumor assessment during therapy.

## RESULTS

### 1. Study overview and tumor-selective EV profiling workflow

We sought to address two key questions: (i) Do tumor-derived EVs carry molecular signals that reflect diagnostic state and therapeutic response in H3K27M-DMG? And (ii) Can these signals be measured using a rapid, multiplex workflow suitable for longitudinal clinical use? To answer these questions, we first evaluated the biological fidelity of EV mRNA signatures using patient-derived DMG cell lines and then translated these findings to clinical plasma specimens. Plasma samples were obtained from 10 patients with H3K27M-DMG and 20 non-DMG healthy controls (n = 30 individuals; 40 total plasma samples). For treatment-response analyses, serial samples were collected from a subset of patients receiving ONC201 **(Fig. 1a)**. Our strategy was designed to complement existing modalities, including MRI, by providing minimally invasive molecular measurements that can be performed repeatedly over time. To enable tumor-selective profiling of circulating EVs, we developed ExPLEX, which integrates immuno-enrichment of DMG-derived EVs with multiplex RNA detection in a same-day workflow **(Fig. 1b)**. In this proof-of-concept study, we applied a focused 4–6 marker panel to evaluate biological relevance and clinical feasibility.

**Figure 1.**
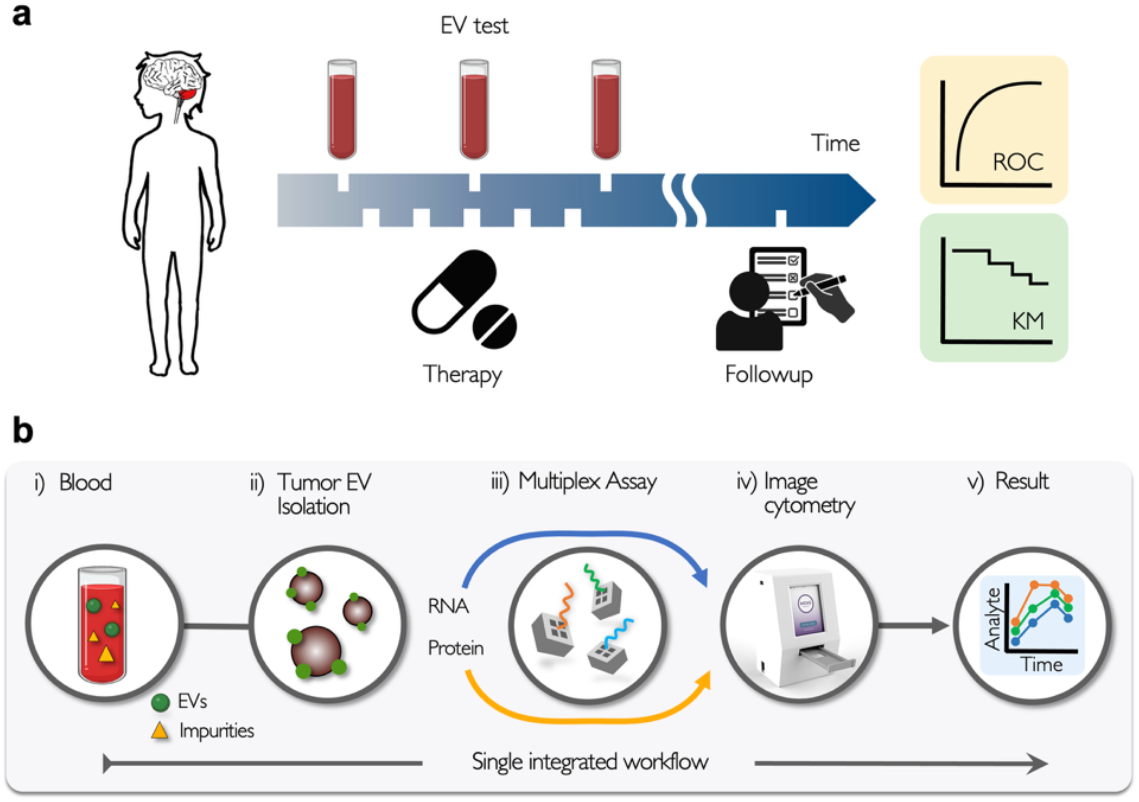
Overview of EV-based molecular monitoring in pediatric DMG. **(a)** Schematic of longitudinal blood-based profiling of tumor-derived EVs for assessment of disease status and therapeutic response. **(b)** Workflow for tumor-selective EV enrichment from plasma followed by multiplex RNA analysis.

### 2. EV mRNA profiles recapitulate parental DMG transcriptomes

We first characterized EVs derived from DMG cell lines to confirm their identity and suitability for downstream analysis. Transmission electron microscopy (TEM) confirmed intact vesicles with characteristic morphology and diameters of ∼50–200 nm, with a mean diameter of ∼150 nm, consistent with small EVs or exosomes (**Fig. 2a**). Complementary physicochemical characterization showed a narrow hydrodynamic size distribution, characteristic negative zeta potential, and minimal protein contamination, supporting the purity and quality of the isolated DMG-EVs (**Supplementary Fig. 1**). Immunoblotting further verified enrichment of generic EV markers, including CD9, CD63, CD81, and TSG101 (**Fig. 2b**).^35^

**Figure 2.**
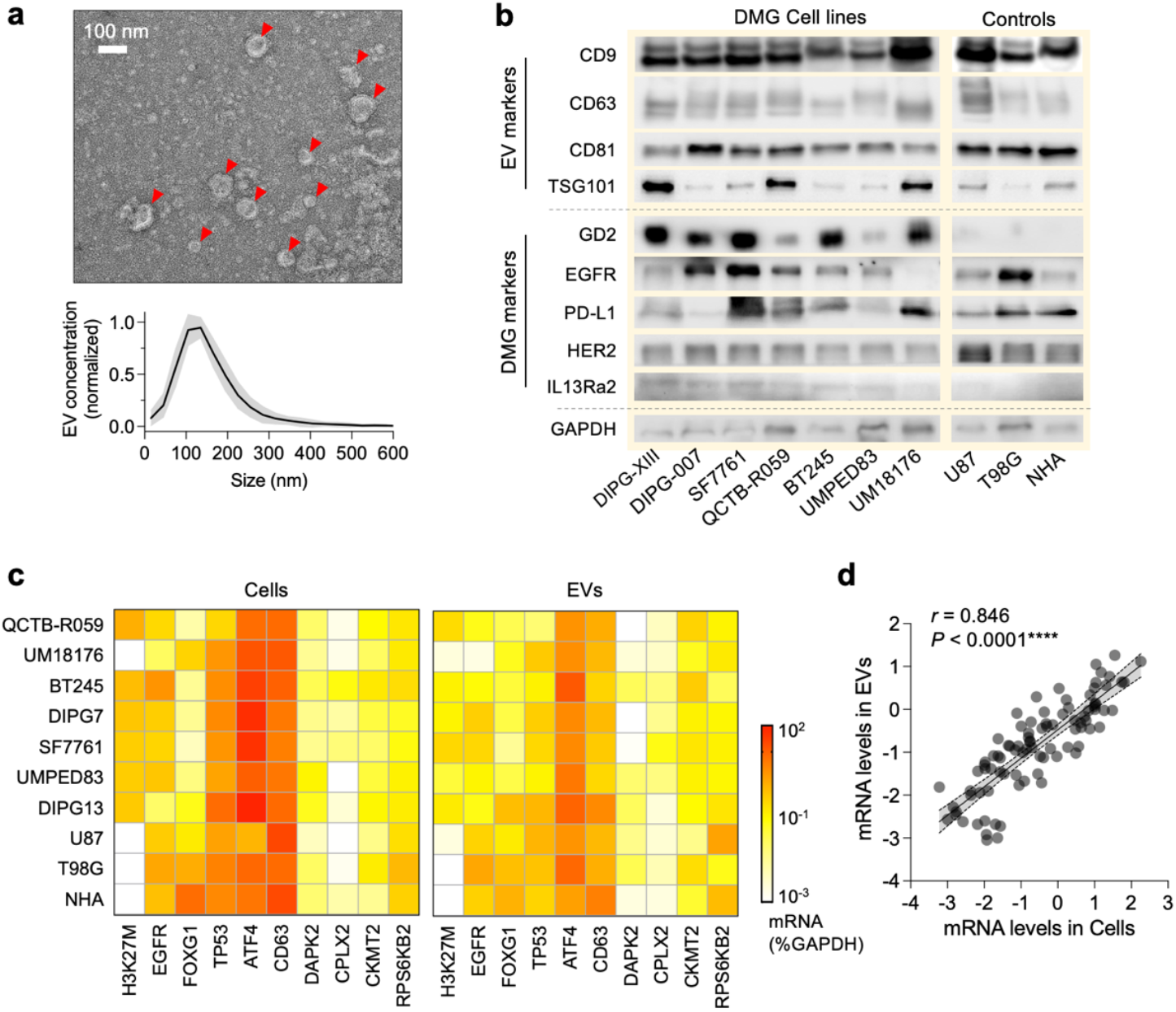
EV protein and mRNA analysis in DMG cell lines. **(a)** Transmission electron microscopy (TEM) and nanoparticle tracking analysis showing EV morphology and size distribution (∼50–200 nm). **(b)** Western blot analysis of EV protein markers from ten DMG cell lines for generic EV markers (CD9, CD63, CD81, TSG101) and tumor-associated markers (GD2, EGFR, PD-L1, IL13Rα2, HER2). **(c)** Heatmap of mRNA levels in parental DMG cells (left) and corresponding EVs (right). Markers include EGFR, FOXG1, ATF4, TP53, and CD63. Expression levels were normalized to GAPDH. **(d)** Correlation between cellular and EV mRNA levels across markers and cell lines (Pearson correlation). Data are shown as mean values from at least three independent experiments.

We then examined whether EV mRNA profiles faithfully reflect the transcriptomic state of their parental DMG cells. To address this, we analyzed a panel of transcripts spanning four functional categories: (i) the DMG diagnostic marker H3F3A (H3K27M); (ii) genes associated with ONC201 treatment response (EGFR, FOXG1, DAPK2, CPLX2, CKMT2, RPS6KB2); (iii) genes involved in cellular stress response (TP53, ATF4); and (iv) a generic EV marker (CD63). We analyzed a panel of human DMG cell lines, with glioblastoma (GBM) cell lines and NHA included as controls. Cellular mRNA levels were quantified by qPCR, and matched EV-derived mRNA levels were measured from EVs isolated from corresponding conditioned media. Across transcripts and cell types, heatmap analysis revealed that relative expression patterns were largely preserved between parental cells and their EVs (**Fig. 2c**). Consistent with this observation, quantitative comparison showed a strong correlation between cellular and EV-derived mRNA levels (Pearson *r* = 0.846, *P* < 0.0001; **Fig. 2d**). Although partial discordance was observed for some low-abundance transcripts, this is biologically plausible because EV cargo does not simply reflect a random sampling of total cellular RNA, and low-input EV RNA may introduce variability during pre-amplification. Therefore, isolated low-level discordant signals were interpreted cautiously and were not considered definitive evidence of transcript presence. These results indicate that EVs retain transcriptomic signatures reflective of the intracellular molecular state of DMG cells, supporting EV mRNA as a circulating readout of tumor state.

### 3. Tumor-selective enrichment of DMG-derived EVs

Because circulating EVs are heterogeneous—originating from both tumor and host tissues— selective enrichment is essential for extracting tumor-specific molecular information from plasma. To this end, we developed a tumor-selective enrichment strategy by screening a panel of EV markers reported to be enriched in DMG or broadly associated with cancer biology.

Candidate markers were selected based on their relevance to glioma, membrane accessibility, and compatibility with immunoaffinity capture. We evaluated their expression across a panel of EVs derived from seven DMG cell lines (DIPG-XIII, DIPG-007, SF7761, QCTB-R059, BT245, UMPED83, and UM18176), two GBM cell lines (U87 and T98G), and normal human astrocytes (NHA). Among the markers tested (GD2, IL13Rα2, EGFR, PD-L1, and HER2), GD2, EGFR, and IL13Rα2 showed preferential expression on DMG-derived EVs compared with EVs from NHA, whereas PD-L1 and HER2 did not (**Fig. 2b**). Based on these results, we combined GD2, EGFR, and IL-13Rα2 into a pooled capture panel (“DMG-pan”) for subsequent immunomagnetic enrichment.

We next evaluated the efficiency and specificity of DMG-pan-based immuno-enrichment using EVs derived from H3K27M-DMG cells. Overall, DMG-pan-functionalized magnetic microbeads achieved high capture efficiency of DMG-derived EVs (∼84%), outperforming conventional tetraspanin-based microbeads (CD-pan; CD9/CD63/CD81; ∼67%) (**Fig. 3a**). Off-target binding was <2%, as determined using microbeads functionalized with IgG control antibodies, indicating a minimal non-specific binding on microbeads (**Fig. 3a** and **Supplementary Fig. 2**). Importantly, whereas CD-pan microbeads captured EVs from both DMG and NHA cells with similar efficiency, DMG-pan beads selectively enriched DMG-derived EVs (∼70%) while exhibiting minimal binding to NHA EVs (∼18%). This minimal binding was statistically indistinguishable from the off-target binding observed with IgG control beads (*P* = 0.1373, two-sided t-test), demonstrating both high efficiency and tumor specificity (**Fig. 3b**).

**Figure 3.**
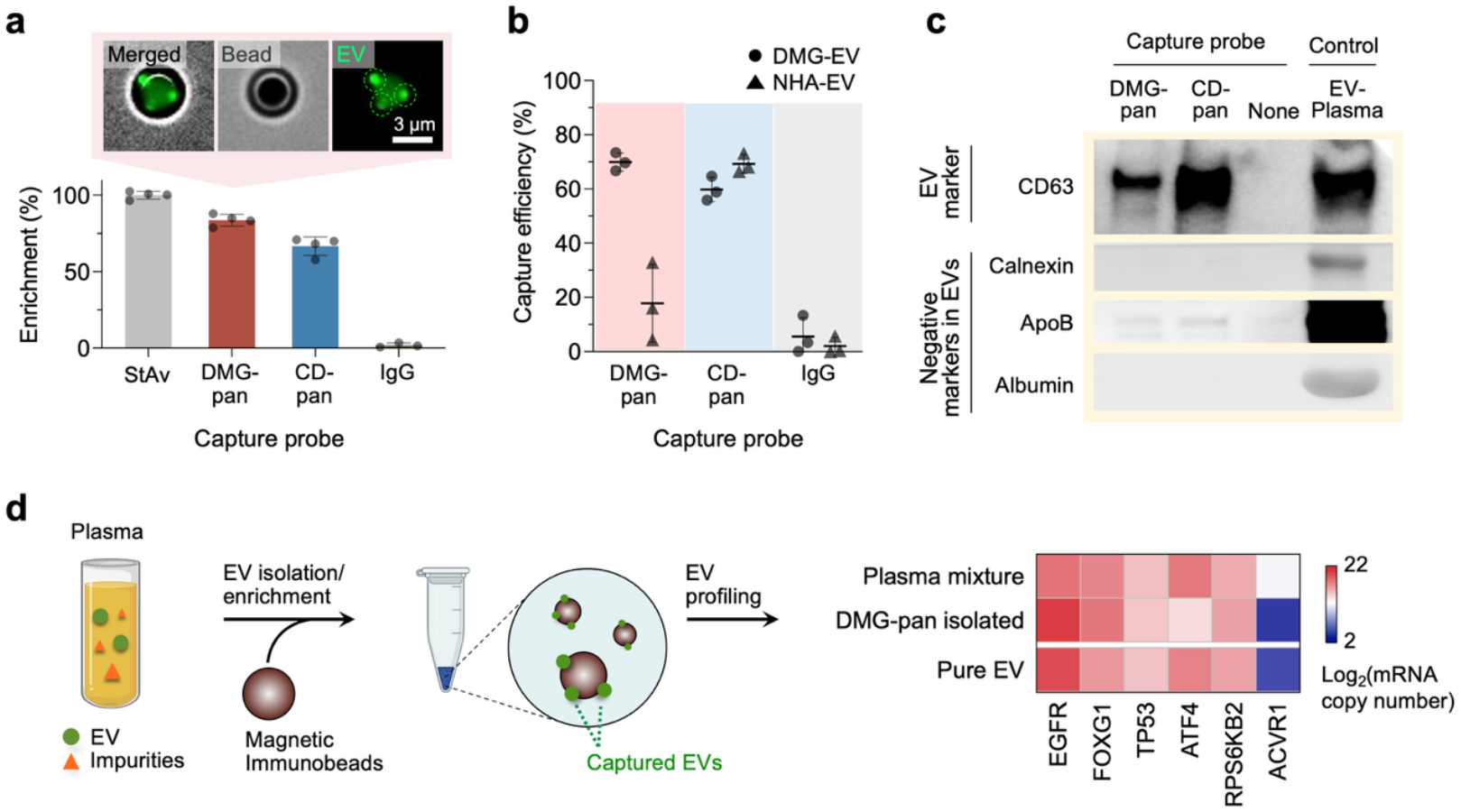
Selective enrichment of DMG-derived EVs. **(a)** Capture efficiency of magnetic immunobeads functionalized with anti–DMG-pan (EGFR, GD2, IL13Rα2), anti–CD-pan (CD63, CD9, CD81), or IgG control antibodies. Values were normalized to streptavidin (StAv)–biotin controls. **(b)** Enrichment specificity using EVs from BT245 DMG cells and normal human astrocytes (NHA). **(c)** Enrichment from EV-spiked human plasma. Immunoblot analysis of enriched fractions showing EV marker CD63 and plasma proteins (Calnexin, ApoB, Albumin). **(d)** mRNA profiles of DMG-pan–captured EVs compared with purified EVs and unenriched plasma EV mixtures. Data are shown as mean ± s.d. (b,c) or mean (d) from at least three independent experiments.

Finally, we assessed enrichment performance in plasma, where background complexity poses the greatest challenge. In spiked plasma samples, DMG-pan enrichment preserved EV markers (CD63) while substantially depleting non-EV plasma proteins, including calnexin, ApoB, and albumin (**Fig. 3c**). This selective enrichment directly improved downstream molecular analysis: mRNA profiles obtained from DMG-pan-isolated EVs closely resembled those of purified DMG EVs and were clearly distinct from unenriched plasma EV mixtures (**Fig. 3d**).

### 4. EV mRNA markers are associated with ONC201 response in vitro

Having shown that EV mRNA profiles recapitulate cellular transcriptomes, we next asked whether EV mRNA markers are associated with therapeutic response to ONC201. EGFR was selected based on prior clinical and preclinical studies identifying EGFR as a molecular determinant of response to ONC201 in GBM, where higher EGFR expression was associated with reduced therapeutic response.^36^ Given the clinical relevance of ONC201 in H3K27M-mutant DMG, including recent FDA accelerated approval for recurrent disease, we sought to test whether this EGFR-associated response pattern could also be observed in this disease context. FOXG1 was included because prior work showed that EGFR mutation can induce FOXG1 expression and EGFR-linked transcriptional changes in GBM,^37^ and because FOXG1 was also supported by our complementary glioma cell-line analyses and baseline tumor RNA-seq data from ONC201-treated H3K27M-mutant DMG patients.^15^ Although FOXG1 is linked to forebrain/cortical developmental features, our analysis of genetically engineered H3K27M-mutant DMG models derived from the same parental line showed that FOXG1 expression can vary within a shared DMG cellular background (**Supplementary Fig. 3**). We therefore evaluated EGFR and FOXG1 as proof-of-concept EV RNA markers associated with ONC201 response to test whether EV profiling could capture treatment-relevant molecular features in H3K27M-mutant DMG, rather than to establish a mechanistic EGFR/FOXG1 resistance pathway.

To define ONC201 response phenotypes, we treated a panel of DMG and glioma cell lines with increasing doses of ONC201 and generated dose–response curves. The GBM lines (U87, T98G) and NHA were included as reference non-DMG controls for comparative context and were not intended to model DMG biology directly. Based on ED_10_ values, cell lines were classified as ONC201-resistant (DIPG13, U87, T98G, NHA) or ONC201-sensitive (DIPG7, SF7761, QCTB-R059, BT245, UMPED83, UM18176) (**Fig. 4a** and **Supplementary Fig. 4**).Consistent with a possible contribution of EGFR signaling to ONC201 response, pharmacologic inhibition of EGFR enhanced ONC201 sensitivity in DMG cells (**Supplementary Fig. 5**).

**Figure 4.**
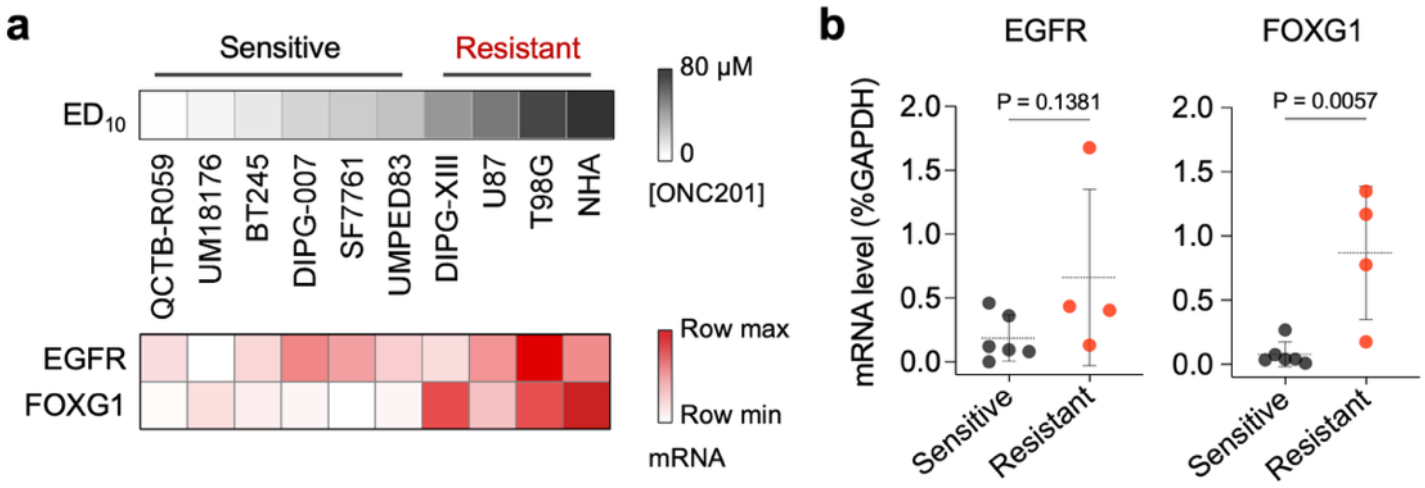
EV mRNA markers associated with ONC201 sensitivity in vitro. **(a)** EV EGFR and FOXG1 mRNA levels in DMG and glioma cell lines plotted against in vitro ONC201 sensitivity (ED_10_). Dose–response curves (top) and corresponding EV mRNA levels (bottom) are shown. “Row maximum” and “row minimum” indicate the highest and lowest EV mRNA expression levels for each marker across cell lines. **(b)** Comparison of EV FOXG1 and EGFR mRNA levels between ONC201-resistant and -sensitive cell lines. Data are shown as mean (a) or mean ± s.d. (b) from at least three independent experiments. ED_10_, 10% effective dose.

EVs were isolated from conditioned media, and EV EGFR and FOXG1 mRNA levels were quantified. Both transcripts showed higher mean EV expression in resistant cell lines compared with sensitive lines (**Fig. 4b**). This difference reached statistical significance for FOXG1 (*P* = 0.0057, two-sided *t*-test), but not for EGFR (*P* = 0.1381, two-sided *t*-test), reflecting overlap in EV EGFR levels between phenotypic groups. Given the heterogeneity of the cell-line panel, including differences in tumor origin, these findings should be interpreted as correlative rather than mechanistic.

### 5. ExPLEX enables high-throughput, multiplexed EV mRNA profiling

Although EV-based diagnostics have attracted increasing attention, clinical translation has been hindered by labor-intensive sample preparation, limited multiplexing capacity, and reliance on specialized instrumentation. To address these barriers, we developed ExPLEX (Extracellular vesicle Multi-PLEXer), a platform designed for scalable, multiplex EV RNA profiling within clinically practical workflows.

ExPLEX integrates three components: (i) shape-encoded hydrogel particle arrays for multiplex target identification; (ii) fluorescence imaging for signal acquisition; and (iii) deep learning (DL) for automated decoding and fluorescence quantification. Each hydrogel particle serves both as an assay substrate and as a physical barcode. For this study, we designed a dot-based encoding scheme in which each particle contains six addressable binary regions, and the presence or absence of a square dot defines a 0/1 pattern, together with an orientation guide for unambiguous code assignment (**Fig. 5a**). This design supports up to 2^6^ = 64 distinct particle identities using a single fluorescence channel, thereby increasing multiplexing capacity through particle shape rather than spectral separation. During readout, paired bright-field (BFM) and fluorescence (FLM) images are acquired in a single scan: the BFM image is used to assign particle identity, and the matched FLM image is used to quantify target-dependent signal from that same particle (**Fig. 5b**). Automated stage scanning and batch DL processing further support scalable analysis across large particle populations (**Supplementary Fig. 6**).

**Figure 5.**
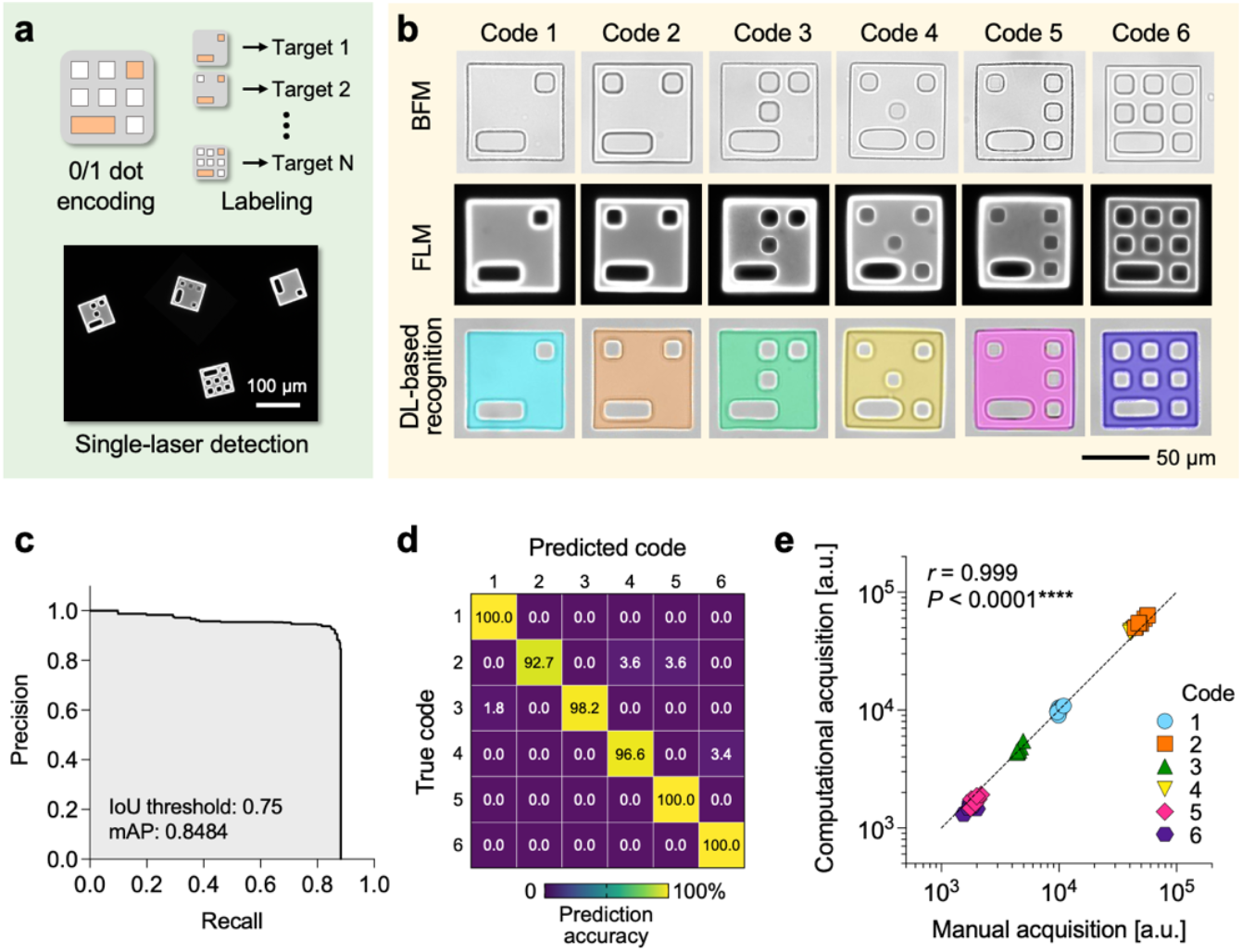
ExPLEX platform and deep learning (DL)-based decoding. **(a)** Schematic of shape-encoded hydrogel particles used for multiplex detection. Each particle contains a binary geometric code defined by the presence (1) or absence (0) of square dot features (white), together with an orientation guide (orange) for unambiguous decoding. The current 6-dot scheme enables up to 2^6^ = 64 unique codes using a single fluorescence channel. **(b)** Representative images of six particle codes acquired by paired bright-field and fluorescence imaging. Top, bright-field (BFM) images used for code recognition; middle, fluorescence (FLM) images acquired after target labeling and used for fluorescence quantification; bottom, false-color DL outputs showing particle detection and code classification from the BFM images. **(c)** Precision–recall curve for the Mask R-CNN–based particle detection model. **(d)** Confusion matrix showing code-classification accuracy across particle types. **(e)** Correlation between DL-quantified and manually measured fluorescence intensities (Pearson correlation). Data are plotted on log–log scales; the dashed line indicates the best-fit linear regression. a.u., arbitrary units.

To ensure robust and unbiased signal extraction, we implemented a Mask R-CNN–based image analysis pipeline. The DL model detects particles and decodes shape codes from the BFM images, and the corresponding FLM images are used to quantify fluorescence intensity at single-particle resolution, as illustrated by the false-color outputs in the bottom row of **Fig. 5b**. At an intersection-over-union (IoU) threshold of 0.75, the network achieved a mean average precision (mAP) of 0.8484 (**Fig. 5c** and **Supplementary Fig. 7**). Code classification accuracies exceeded 92% across particle types (**Fig. 5d**). DL-derived fluorescence measurements showed excellent agreement with manual quantification across a broad dynamic range (Pearson *r* = 0.999, *P* < 0.0001; **Fig. 5e**).

We next evaluated analytical performance for EV mRNA detection. Target-specific capture and detection probes were designed to hybridize defined RNA sequences, followed by fluorescent reporter binding for optical readout (**Fig. 6a**). Serial dilution experiments using synthetic DNA oligonucleotides that mimic EV-derived amplicons demonstrated dose-dependent responses with a limit of detection (LOD) of ∼2.9 pM for FOXG1 (**Fig. 6b,c**). Comparable sensitivity and dynamic range were observed when applied to EV-derived mRNA from DMG cell lines, indicating consistent performance in biologically relevant samples (**Fig. 6c**). Multiplex specificity was assessed across DMG-relevant transcripts, including EGFR, FOXG1, TP53, H3K27M, and GAPDH. ExPLEX exhibited minimal cross-reactivity and robust target discrimination in multiplex format (**Fig. 6d**). Quantitative comparison with qPCR demonstrated strong concordance across markers and cell lines (Pearson *r* = 0.8326, *P* < 0.0001; **Fig. 6e,f**).

**Figure 6.**
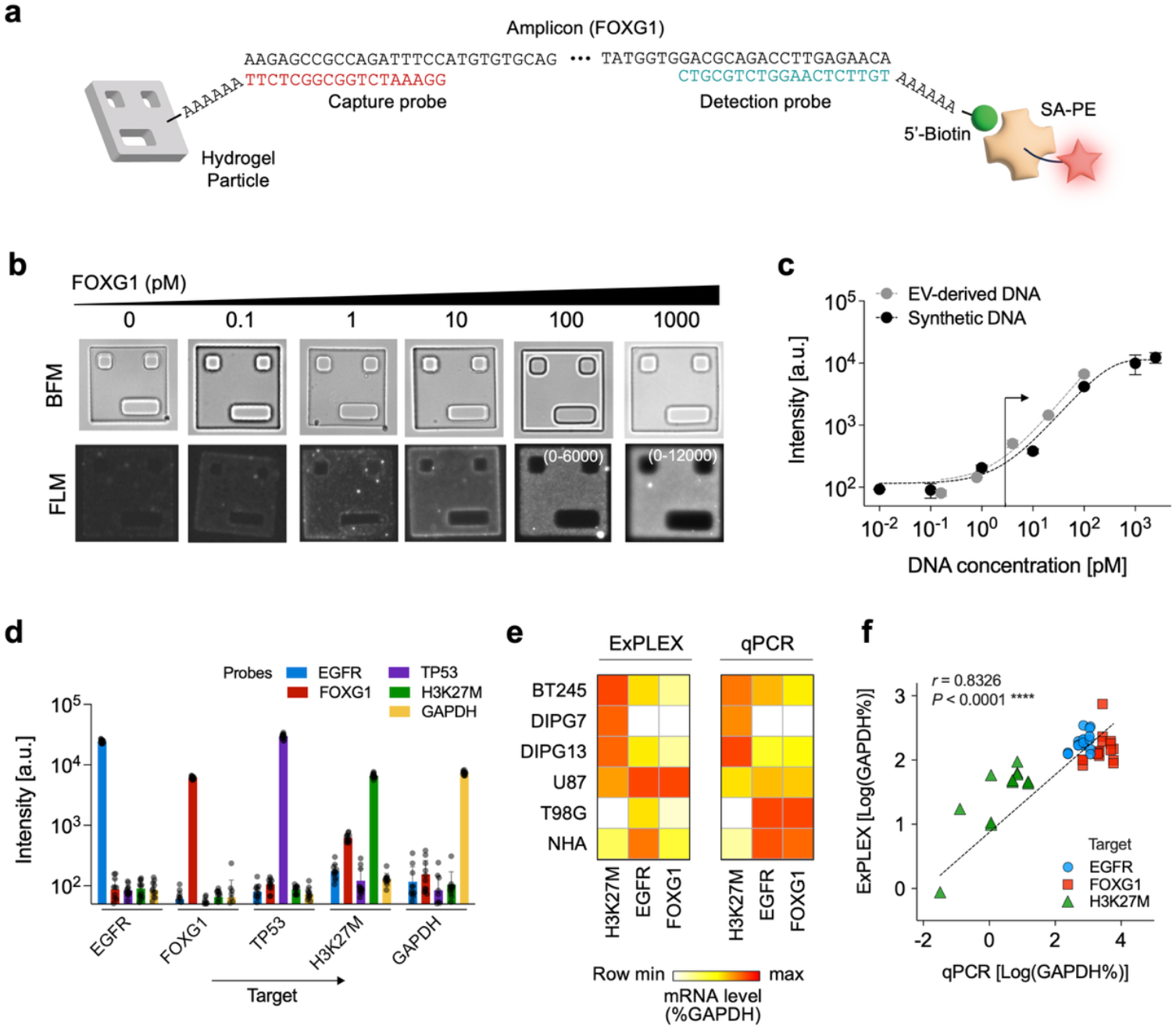
Multiplexed EV mRNA detection using ExPLEX. **(a)** Design of capture and detection probes targeting EV-derived RNA sequences. Two probes differing by a single nucleotide were used for sequence-specific hybridization. **(b)** Representative bright-field (BFM) and fluorescence (FLM) images of hydrogel particles following hybridization with synthetic DNA targets at varying concentrations. **(c)** Analytical sensitivity determined using serial dilutions of synthetic DNA or EV-derived amplicons generated by asymmetric RT-PCR (35 cycles). Synthetic DNA sequences matched EV-derived targets. The limit of detection (LOD) was ∼2.9 pM for FOXG1. **(d)** Differential detection of multiple RNA targets demonstrating high specificity and low cross-reactivity in multiplex format. **(e)** Heatmap comparison of ExPLEX and qPCR measurements across markers and cell lines. **(f)** Correlation between ExPLEX and qPCR quantification across samples (Pearson *r =* 0.8326, *P* < 0.0001). Data are shown on log–log scales; the dashed line indicates the linear regression fit. Data represent mean ± s.d. (c,d) or mean (e,f) from at least three independent experiments. a.u., arbitrary units.

### 6. Clinical validation of EV mRNA markers in pediatric DMG

We next evaluated the clinical performance of EV mRNA markers measured by ExPLEX in plasma samples (and CSF when available) from patients with H3K27M-mutant DMG and non-DMG healthy controls (**Supplementary Table 1**). Specifically, we examined whether EV H3K27M mRNA enables diagnostic discrimination and whether EV EGFR and FOXG1 mRNA levels are associated with clinical outcome following ONC201 therapy.

EV H3K27M mRNA levels were higher in plasma-derived EVs from DMG patients (n = 10) compared with non-DMG healthy controls (n = 20) (**Fig. 7a-b**, *P* < 0.0001, two-sided *t*-test). Some background signal was also observed in non-DMG controls, as expected given the single-nucleotide difference between H3K27M and wild-type H3F3A and the low abundance of EV RNA, which warrants threshold-based interpretation.^38^ We therefore defined H3K27M positivity using a preliminary threshold of mean + 3 × SD of the non-DMG control group. We also used 5-fold cross-validation to assess the robustness of this approach, which showed limited variation in the threshold across folds in this cohort (**Supplementary Fig. 8**). Even with this conservative cutoff, the H3K27M signal in DMG patient samples remained clearly elevated relative to non-DMG controls. ROC analysis demonstrated strong separation in this cohort (AUC = 1.00; **Fig. 7c**). In contrast, qPCR analysis of matched samples yielded lower diagnostic performance (AUC = 0.855; **Fig. 7c** and **Supplementary Fig. 9**), indicating improved discrimination using the ExPLEX workflow. Although limited by cohort size, these findings support circulating EV H3K27M mRNA as a minimally invasive diagnostic marker in DMG.

**Figure 7.**
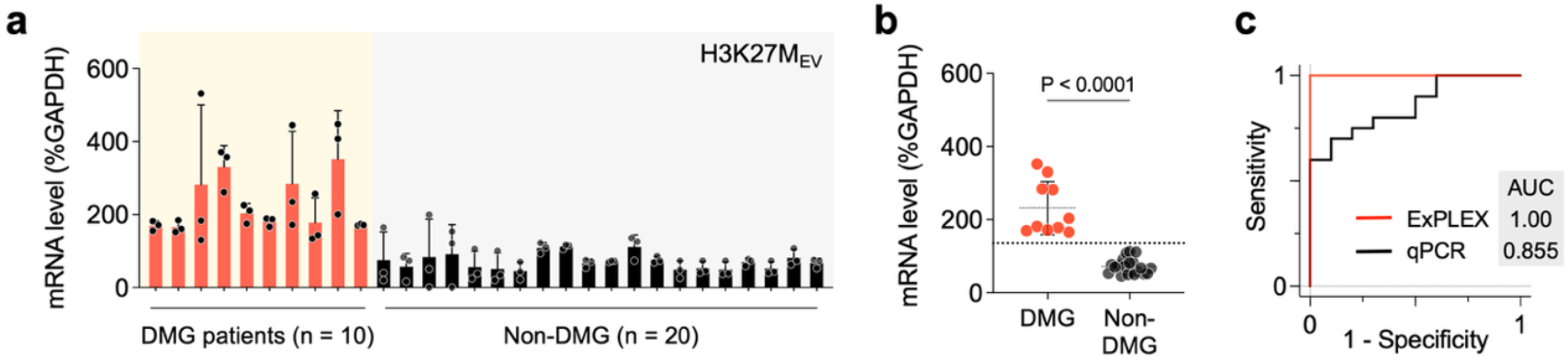
Diagnostic discrimination of DMG using EV mRNA. **(a)** Plasma EV H3K27M mRNA levels measured by ExPLEX in DMG patients and non-DMG healthy controls. **(b)** EV H3K27M mRNA levels were higher in DMG patients than in non-DMG controls (two-sided *t*-test). Dotted line indicates a positive cutoff for H3K27M. **(c)** Receiver operating characteristic (ROC) analysis for classification of DMG vs. non-DMG samples. The area under the curve (AUC) was calculated for ExPLEX and qPCR on matched samples. Data represent mean ± s.d. from three independent measurements.

We next examined, in an exploratory analysis, whether EV EGFR and FOXG1 mRNA levels were associated with clinical outcomes following ONC201 therapy. Plasma samples used in this analysis were collected from ten patients at pre- and post-treatment time points relative to ONC201 therapy. Patients were stratified as long-term survivors (LTS) or short-term survivors (STS) based on progression-free survival (median PFS, 310.5 days). EV EGFR and FOXG1 levels showed substantial inter-patient variability (**Fig. 8a**). When stratified by outcome, EV EGFR and FOXG1 levels were lower in samples from LTS patients than in those from STS patients (**Fig. 8b**; *P* = 0.0029 for EGFR; *P* = 0.0021 for FOXG1). Higher EV EGFR and FOXG1 levels also trended with poorer radiographic response, reflected by greater increases in tumor area relative to baseline (**Fig. 8c**), and with shorter PFS (**Fig. 8d**). ROC analysis showed separation between LTS and STS within this cohort (AUC = 0.88 for EGFR; AUC = 0.89 for FOXG1). Notably, transcript levels measured from available tumor tissue did not clearly distinguish LTS from STS (**Supplementary Fig. 10**). Given the modest cohort size, these findings should be interpreted as preliminary and hypothesis-generating.

**Figure 8.**
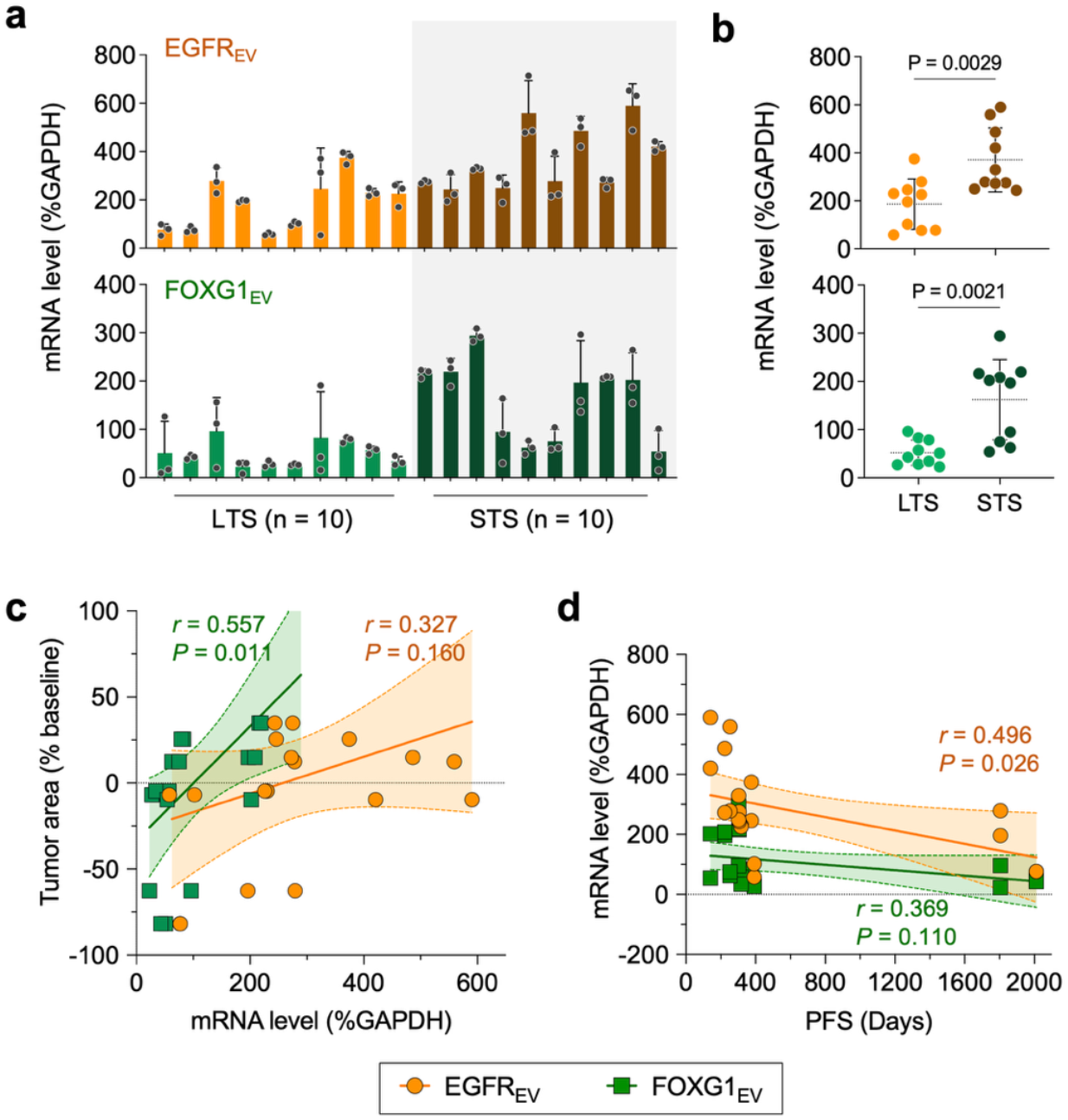
Exploratory association of EV mRNA markers with ONC201 response in H3K27M-DMG. **(a)** EV EGFR and FOXG1 mRNA levels measured in patients receiving ONC201. **(b)** EV EGFR and FOXG1 levels were lower in long-term survivors (LTS, *n* = 5 patients, 10 samples) compared with short-term survivors (STS, *n* = 5 patients, 10 samples) in this exploratory cohort (two-sided t-test). **(c)** Correlation between EV mRNA levels and % tumor area change from baseline (Pearson correlation). **(d)** Correlation between EV mRNA levels and progression-free survival (PFS) (Pearson correlation). Data represent mean ± s.d. from three independent measurements.

## DISCUSSION

Detection of reliable markers of therapy response and drug resistance remains a major clinical challenge in brain cancer. This need is particularly acute in DMG—one of the most lethal pediatric brain tumors—where tumor location precludes resection and serial biopsy, yet treatment response and resistance can rapidly evolve over weeks to months. Liquid biopsy approaches have thus emerged as minimally invasive alternatives to tissue-based monitoring. Recent studies, including our own, have demonstrated the feasibility of cell-free tumor DNAs (ctDNAs)^18,19^ and microRNAs (miRNAs)^20^ in plasma and CSF as minimally invasive DMG biomarkers. However, these approaches face biological and practical constraints, including low analyte abundance in plasma and the logistical complexity and procedural risk associated with repeated CSF sampling. Circulating tumor cells (CTCs) have also been explored,^39^ but they are rare in gliomas and are not routinely detectable even from large blood volumes (∼10 mL), limiting their clinical utility.

EVs provide a compelling complementary liquid biopsy modality for DMG. Tumor-derived EVs are abundant and stable in circulation, readily cross the blood–brain barrier, and can be analyzed from small sample volumes (∼100 *µ*L) of plasma or CSF. Unlike fragmented ctDNA, EVs carry multimodal molecular cargo that reflects active cellular states. These properties motivated our focus on EVs as a vehicle for measuring gene products relevant to therapy response. To date, limited work has examined EVs in pediatric DMG, with prior studies focused primarily on the biological roles of EVs in tumor phenotype modulation and radiation response.^40^ The potential utility of EVs for diagnosis and longitudinal monitoring in H3K27M-DMG has remained largely unexplored. Our findings extend this framework by showing that plasma-derived EV RNA can capture response-associated molecular features in a clinically relevant setting.

Here, we provide evidence that circulating tumor-derived EV mRNA carries potentially clinically useful information in DMG. Specifically, we show that (i) EV mRNA levels correlated strongly with transcript levels in parental DMG tumor cells across multiple targets and cell lines (10 mRNA targets × 10 cell lines, *r* = 0.846), supporting biological fidelity; (ii) EV H3K27M mRNA distinguished patients from non-DMG healthy controls; and (iii) EV EGFR and FOXG1 mRNA levels were associated with radiographic response and PFS following ONC201 therapy in an exploratory cohort. These findings suggest that EV mRNA profiling may provide a dynamic molecular readout that complements conventional imaging and tissue-based assessment.

Translating EV biomarkers into clinical practice requires practical and scalable assays. To address technical barriers, we developed ExPLEX, which integrates tumor-selective immuno-enrichment with multiplex RNA profiling in a same-day workflow. ExPLEX achieved efficient enrichment of DMG-derived EVs, sensitive multiplex detection (LOD ∼2.9 pM for FOXG1), and strong concordance with qPCR (r = 0.8326). Diagnostic discrimination using ExPLEX exceeded that of qPCR in this cohort. By implementing multiplexing through geometric encoding rather than additional fluorophores, the platform enables scalable expansion of biomarker panels without increasing optical complexity, supporting practical clinical deployment.

ExPLEX builds on our prior work in EV profiling and multiplex molecular sensing. Earlier platforms enabled sensitive detection of EV surface proteins using plasmonic^41^ and magneto-electrochemical^42^ approaches, but provided limited access to intravesicular cargo. In parallel, we developed shape encoded-particle-based platforms for multiplexed molecular analysis in other contexts,^43,44^ but those systems lacked integrated EV enrichment and automated decoding. ExPLEX integrates these elements into a single unified workflow, enabling direct interrogation of intravesicular RNA markers relevant to therapy response. The geometric encoding strategy supports panel expansion without additional spectral burden, facilitating practical implementation.

The clinical implications are significant. MRI remains the standard for therapy assessment in DMG but primarily reflects anatomical changes and may be confounded by treatment-related effects such as pseudo-progression. EV mRNA profiling offers a complementary molecular perspective on tumor state during therapy. By capturing transcriptional features associated with treatment response, EV profiling may provide molecular information that complements imaging. More broadly, EV-based profiling can be expanded to incorporate additional resistance-associated transcripts and integrated with mutation-based liquid biopsy assays for longitudinal monitoring. Beyond DMG, the modular design of ExPLEX allows adaptation to other tumor types and therapeutic contexts in which serial tissue access is limited. Capture antibodies and probe sets can be reconfigured without altering the core detection workflow, providing a generalizable framework for scalable molecular monitoring across diverse solid tumors. Future studies will determine the extent to which this strategy generalizes across oncologic settings.

As a proof-of-concept study, several limitations need to be addressed. First, the cohort size is modest, and performance estimates should be considered exploratory and hypothesis-generating. In addition, the survival-associated EV analysis was performed in a relatively small cohort without multivariable adjustment for known clinical and molecular prognostic factors, and sample timing was not fully uniform across patients. These findings should therefore be considered preliminary, and larger, multi-institutional studies with standardized sampling schedules will be required to establish robust thresholds and assess generalizability. Second, plasma EV H3K27M levels may be influenced by biological factors beyond tumor burden alone, including blood–brain barrier integrity. Accordingly, these measurements should be interpreted as complementary rather than standalone biomarkers, and their clinical utility will require validation in larger prospective cohorts. Third, FOXG1 expression is influenced by regional tumor context, which may contribute to its association with ONC201 response, although our analyses in isogenic DMG models suggest that this association is not solely explained by tumor origin. Fourth, systematic head-to-head comparisons with ctDNA in matched plasma and CSF samples—particularly in clinically challenging scenarios such as pseudo-progression—will be important to delineate where EV profiling provides unique or complementary value. Finally, this study demonstrated modest multiplexing (5-plex detection using a 64-code particle library) as an initial feasibility test. ExPLEX is not intended to compete with high-end platforms such as spectral flow or mass cytometry, but rather to provide a compact and accessible system for scalable EV molecular profiling in near-patient settings. Future work will focus on expanding multiplexing capacity, incorporating additional analyte classes (including EV proteins), and advancing toward a fully automated sample-in, answer-out format through integration with microfluidic cartridges^45,46^ and simplified sample-prep workflows.^47-49^

In summary, this work supports tumor-derived EV mRNA as a biologically informative and potentially clinically useful biomarker modality in pediatric DMG and introduces ExPLEX as an enabling platform for tumor-selective, multiplexed EV RNA profiling from blood within a same-day workflow. By coupling disease-focused EV enrichment with scalable multiplexing and automated decoding, this approach supports both the biological rationale and the practical feasibility of EV-based molecular monitoring in clinical oncology.

## METHODS

### 1. Cell culture

All cell lines and culture conditions are summarized in **Supplementary Tables 2 and 3**. Human diffuse midline glioma (DMG) cell lines were cultured as spheroids in tumor stem media (TSM) at 37 °C under 5% CO_2_, initiated at 2.5 × 10^5^ cells/mL. Spheroids were passaged upon reaching several hundred micrometers by centrifugation (350 × g, 4 min) followed by Accutase dissociation. Glioblastoma (GBM) cell lines and normal human astrocytes (NHA) were cultured in DMEM and passaged every 4 days using 0.05% Trypsin-EDTA. NHA cultures were used at passages 3–5. Conditioned media were collected, centrifuged at 350 × g to remove cells and at 3,000 × g for 30 min to remove debris, and stored at −80 °C until EV isolation. Three independent cultures were used per cell line. All cell lines were routinely tested and confirmed to be free of mycoplasma contamination using two independent assays.

### 2. In vitro drug treatment

ONC201 was used to evaluate treatment response in DMG and glioma cell lines. To determine drug sensitivity, cells were seeded at a density of 5 × 10^3^ cells per well in 96-well plates and allowed to stabilize overnight. Cells were then treated with ONC201 at final concentrations ranging from 0.1 to 100 μM for 72 h. Cell viability was assessed using CellTiter-Glo (Promega) or CCK-8 (Abcam) assays according to the manufacturers’ protocols. Luminescence or absorbance signals were measured using a microplate reader (Synergy H1, BioTek).For downstream EV and molecular analyses, cells were treated with ONC201 at concentrations corresponding to their respective effective doses, as determined from viability assays.

### 3. EV isolation and characterization

The frozen media was equilibrated at room temperature and filtered with 0.4 µm syringe filter. The filtrates were concentrated with a centrifugal filter (Merck, Amicon-70, MWCO=10K) into 1 mL of volume and loaded in the size exclusive chromatography (SEC) column (Cytiva, Sepharose CL-4B). In the 10 mL of SEC column, loaded EV concentrate was eluted with PBS, obtaining a 2.5 mL of first and second-EV fraction, respectively. The EV fraction was further concentrated with a centrifugal filter (Merck, Amicon-2) into 0.2 mL of EV solution. For the measurement of EV size distribution and concentration, nanoparticle tracking analysis (NTA, Particle Metrix Inc., ZetaView) and bicinchoninic acid assay (BCA assay, Thermofisher) were performed, followed by storing the sample at -80 °C before use. The EV samples were lysated with RIPA buffer for 30 min in the ice bath and centrifuged at 16,000 x g for 15 min to analyze the EV protein. For EV characterization, the size distribution and zeta potential of isolated EV were measured using dynamic light scattering (DLS) spectrometer (Zetasizer Nano ZS, Malvern Panalytical Inc.) and nanoparticle tracking analysis (NTA). The size and morphology of DMG derived EVs were characterized with a transmission electron microscope (Talos F200X, ThermoFisher). The EV solution was diluted with DI water and dropped onto the copper-carbon grid (Ted Pella Inc.), followed by drying in a vacuum chamber.

### 4. EV protein and lipid analysis

#### Immuno-blotting analysis for EV protein

Antibodies used for immunoblotting are listed in **Supplementary Table 4**. Quantified EVs samples were lysed in the RIPA buffer containing protease inhibitors (Cell Signaling Technology) and incubated in the ice bath for 30 min. The lysed EV samples were centrifuged at 16,000 x g for 15 minutes to remove cell debris and denatured using sample buffer (Invitrogen, LDS sample buffer) at 95 °C for 5 minutes. The protein lysates were developed by sodium dodecyl sulfate-polyacrylamide gel electrophoresis (SDS-PAGE) at 150 V for 40 min and transferred onto the nitrocellulose membrane (Bio-Rad) at 80 V for 90 min. The nitrocellulose membrane was blocked with 5% skim milk-TBST buffer for 1 hour and the blocked membranes were immune-blotted with various antibodies at 4 °C overnight according to the described concentration. Following incubation with HRP-conjugated secondary antibodies at room temperature for 2 hours, sequential washing process, chemiluminescence reagent (Thermofisher, Supersignal Pico, femto & atto) was applied on the nitrocellulose membrane for several minutes. The chemiluminescence signal of the nitrocellulose membranes were imaged through a gel imaging system (ChemiDoc, Bio-Rad).

#### Immuno-blotting analysis for lipid

For the GD2 glycolipid detection, immuno-stanning through thin layer chromatography (TLC) assay was performed. The 20 *µ*L of EV samples were mixed with 300 *µ*L of chloroform-methanol solution (2:1, (v/v)) and 40 *µ*L of DI water was added. Mixed solutions were centrifuged at 15,000 x g for 3 minutes to separate the organic and aqueous phase. The bottom of the organic phase was transferred into a new tube and dried the solvent to concentrate the lipid. Dried lipid pellets were re-dissolved with a small volume of chloroform and deposited onto the TLC aluminum sheet (Merck, TLC silica gel 60 F_254_) with a glass capillary. The lipid samples on the TLC membrane were developed in the mobile phase [Chloroform: Methanol: 0.5 wt% CaCl_2_ water = 55:45:10, (v/v)] and immunoblotted in the same manner as nitrocellulose membranes. Chemiluminescence reagent was applied onto the TLC plate and chemiluminescence signal of the TLC plates were imaged through a gel imaging system (ChemiDoc, Bio-Rad).

### 5. Immunomagnetic enrichment and EV labeling

#### Magnetic beads modification

Antibodies used for immunomagnetic separation are listed in **Supplementary Table 4**. Epoxy-modified magnetic beads [1.0 mg] (Dynabeads™ M-270 Epoxy, Invitrogen) were washed with PBS for 10 minutes and sedimented within a magnetic rack. Single or pooled antibody solution [25 μg of protein] was added to the magnetic beads and mixed in the hula mixer for 24 hours at 37 °C. Antibody conjugated magnetic beads were washed with 0.05% tween-20 in the PBS buffer twice and blocked with 0.1% of BSA in the PBS solution for 2 hours at room temperature. Antibody-functionalized magnetic beads were further washed with PBS and stored in the 0.1 % BSA-PBS buffer at 4 °C before use.

#### EV modification

Fluorescence EVs were prepared with PKH dyes (PKH26 and PKH67 fluorescence cell linker, Merck) through the manufacturer’s protocol. EV samples were diluted with diluent solution and 6 *µ*L of dye solution was added for 1 mL of sample volume. Mixed solutions were incubated at room temperature for 5 minutes and quenched with 2 mL of 10 wt% BSA-PBS solution. Add PBS buffer to reach 8.5 mL of total volume and transfer into an ultracentrifuge tube. 1.5 mL of sucrose solution (0.971M) was slowly embedded on the bottom of the tube and centrifuged at 190,000 x g for 2 hours at 4 °C. The supernatant was removed and EV pellet was resuspended in PBS, followed by washing EVs as the same conditions. For preparing biotinylated EVs, EV samples were treated with 5.0 mM concentration of Sulfo-NHS-LC-biotin compound (EZ link, Thermofisher) and incubated the solution for 1 hour under stirring. The prepared sample was diluted with PBS and concentrated with Amicon-2, resulting in 100 μL of biotinylated EVs with 88.6 % of yield.

### 6. EV RNA extraction and analysis

#### EV RNA extraction and quantification

For the RNA extraction of cells, cell pellets (suspension cells) or cells on the bottom of culture flasks (adhesion cells) were treated with TRIzol reagent (Invitrogen). The lysate solutions were transferred to the tube and mixed well to get the complete lysate. To separate the phases, chloroform was mixed into the tube and incubated in an ice bath for 5 minutes. The solution was centrifuged at 16,000 x g for 10 minutes and the aqueous phase was transferred into a new tube. The same amount of IPA was added into the tube and centrifuge the solution at 16,000 x g for 10 minutes to make RNA pellets. The RNA pellets were washed with 75% (v/v) ethanol twice and dried under the hood, resulting in a transparent RNA pellet. The RNA pellets were dissolved in the RNase-free water and quantified using a Nanodrop spectrometer (Thermo scientific). For the RNA extraction of EVs, a silica column kit (Total Exosome RNA and Protein isolation kit, Invitrogen or Directzol RNA microprep, Zymo research) was used according to the manufacturer’s protocol. Isolated RNA solutions were quantified with Bioanalyzer 2100 (Agilent) using RNA Pico chips for the downstream analysis. All RNA samples were immediately used for downstream process or stored at -80 °C before use.

#### EV mRNA analysis via Qpcr

Extracted RNA was converted into cDNA using reverse-transcription kit (High-Capacity cDNA Reverse Transcription, Applied Biosystems). EV derived cDNA samples were pre-amplified (Taqman PreAmp Master Mix, Life Technologies) through 14 cycles before qPCR. The qPCR reactions were performed using Powerup SYBR green Master Mix or Taqman Master mix (Life Technologies) and 500 nM of each primer set according to the manufacturer’s protocol. The qPCR cycle consisted of 1 cycle of 50 °C for 2 min and 95 °C for 2 min, 40 cycles of 95 °C for 15 sec, 57 °C for 15 sec, and 72 °C for 1 min. The conventional qPCR was performed with QuantStudio 3 (Applied Biosystems). All qPCR experiments were done in triplicate and relative gene expression was calculated and normalized by respective GAPDH expression.

### 7. Target amplification for ExPLEX analysis

#### EV isolation and mRNA extraction

EV samples dispersed in PBS buffer or plasma were incubated with immuno-magnetic beads for 1 h with standing on hula mixer. The NTA was performed in the supernatant to calculate the remaining EVs. The immuno-magnetic beads were washed with PBS three times and captured EVs were lysated with Trizol or RIPA buffer according to the downstream analysis. A silica column kit (Total Exosome RNA and Protein isolation kit, Invitrogen or Directzol RNA microprep, Zymo research) was used according to the manufacturer’s protocol. Isolated RNA solutions were quantified with Bioanalyzer 2100 (Agilent) using RNA Pico chips for the downstream analysis. All RNA samples were immediately used for downstream process or stored at -80 °C before use.

#### Amplification of target gene

Extracted RNA was converted into cDNA using reverse-transcription kit (High-Capacity cDNA Reverse Transcription, Applied Biosystems) and EV derived cDNA samples were asymmetric-amplified before ExPLEX assay. The asymmetric PCR were performed using GoTaq G2 Colorless Master Mix (Promega) and 500 nM of forward primer and 50 nM of reverse primer. PCR cycle consisted of 1 cycle of 95 °C for 10 min, 35 cycles of 94 °C for 15 sec, 58 °C for 25 sec, and 72 °C for 30 sec, and 1 cycle of 72 °C for 7 min.

### 8. Hydrogel particle synthesis and functionalization

#### Particle synthesis

Shape-encoded hydrogel microparticles were fabricated via a discontinuous dewetting (DD) technique using a degassed PDMS mold, as previously described with minor modifications (*REF. H*.*U*.*K. 2020*). A master mold with the desired features was first prepared using SU-8 photolithography. PDMS was then cast against the master mold and cured at 70 °C for 4 h to generate a flexible negative mold. Prior to use, the PDMS mold was placed in a vacuum chamber and degassed for 10 min to enable efficient precursor filling via gas-absorption-driven wetting of microwells. A prepolymer solution composed of 20% (v/v) PEGDA-700, 75% (v/v) PEG 600, and 5% (v/v) Darocur 1173 photoinitiator was loaded onto the surface of the degassed mold. After the precursor filled the microwells, the excess precursor on the mold surface was removed by gently scraping with a glass slide. The mold was then exposed to UV light (365 nm, 200 mW/cm^2^) for 145 ms to initiate photopolymerization. Polymerized microparticles were collected by adding PBS containing 0.05% Tween-20 (PBST) onto the PDMS mold and gently pipetting repeatedly until the particles detached from the mold. Collected particles were subsequently washed five times with PBST before the post-functionalization.

#### Functionalization

To introduce probe-specific functionality, single-stranded DNA probes were conjugated to the collected hydrogel microparticles through thiol-ene reaction, as previously reported (*REF*.). Briefly, thiolated DNA probes were first reduced in 0.5 mM TCEP at 24 °C for 1 h. The reduced probes were then incubated with the microparticles at 37 °C for 48 h in a thermal shaker (Model #13687711, Thermo Fisher) with agitation at 1400 rpm. Following conjugation, the particles were rinsed eight times with a TE buffer containing 0.05% Tween-20 (TET) to remove unbound probes.

### 9. ExPLEX Assay for RNA profiling

Sequences for species-specific primers targeting EV RNAs are listed in **Supplementary Table 5**. The ExPLEX assay was performed through sequential steps of target hybridization, biotin conjugation, fluorescent labeling, and imaging. For target hybridization, 10 *µ*L of asymmetric PCR product were heated at 90 °C for 5 min, snap-cooled for 5 min, and mixed with 15 *µ*L of hydrogel microparticles (∼ 50 particles) and 25 *µ*L of 1× TET buffer containing 500 mM NaCl, to obtain a final NaCl concentration of 250 mM in a total volume of 50 *µ*L. The mixture was incubated at 50 °C for 1 h in a thermal shaker (Model #13687711, Thermo Fisher) at 1,400 rpm. Microparticles were then washed three times with TET buffer containing 50 mM NaCl and resuspended in 35 *µ*L. For biotin conjugation, 8 *µ*L of a 0.1 mM biotin-conjugated probe and 36 *µ*L of TET buffer with 500 mM NaCl were added to the 35 *µ*L of target-hybridized microparticles to maintain a final NaCl concentration of 250 mM. The probe was designed to hybridize to the target mRNA amplicon immobilized on the hydrogel microparticles. The reaction proceeded at 50 °C for 1 h at 1,400 rpm, followed by three washes with TET buffer containing 50 mM NaCl and resuspension in 45 *µ*L. For fluorescent labeling, streptavidin-conjugated phycoerythrin (SA-PE; S866, Invitrogen) diluted to 0.1 mg/mL in PBS was added (5 *µ*L) to the biotin-labeled microparticles and incubated in the dark at room temperature for 30 min at 1,400 rpm. Finally, microparticles were washed three times with TET buffer containing 50 mM NaCl to remove unbound SA-PE, and fluorescence images were acquired using an Axio Imager M2 (Zeiss) microscope equipped with an R-PE filter set (Ex/Em = 565/576 nm) and a 10× objective lens.

### 10. Training and evaluation of deep learning model

The Mask R-CNN model was trained by transfer learning. The network was initialized with COCO-pretrained parameters (trained until 12 epochs, as provided by MMDetection).^50^ Training was performed using stochastic gradient descent (SGD) with a step learning-rate scheduler. The initial learning rate was set to 10-2 and decayed by a factor of 0.1 every 20 epochs. The model was trained for a total of 10 0 epochs, and checkpoints were saved at each epoch. For training data preparation, 168 images were manually annotated using the Supervisely annotation tool. In addition, 336 synthetic images were generated for data augmentation,^51^ resulting in a total of 504 images. The dataset was split into training and validation sets at an 8:2 ratio. After training, the checkpoint from epoch 21, which achieved the lowest validation loss, was selected for subsequent analyses. Model performance was further evaluated using an independent test set of 101 images that were not used during training and mean average precision (mAP) and a confusion matrix were evaluated.

### 11. Deep learning-based analysis software

A deep learning–based analysis software was developed in Python with a graphical user interface (GUI) for user-friendly, automated processing. The software automatically loads paired bright-field and fluorescence images from a designated folder, analyzes each image pair, and reports summary statistics (e.g., mean and standard deviation) of fluorescence intensity for each particle code. Image analysis was performed using a Mask R-CNN model applied to the bright-field images to generate pixel-wise masks for individual particles and to classify their intrinsic shape. The masks were used to quantify fluorescence intensity from the corresponding fluorescence images, while the shape classification was used to assign particle codes for biomarker identification. For background correction, the fluorescence intensity of each particle was calculated after subtracting the mean background intensity measured from particle-free regions within the same image. After basic statistical analysis, the software calculates biomarker concentrations using a predefined calibration curve that correlates fluorescence intensity with target concentration.

### 12. Reference assays for benchmarking

#### ELISA

We used the same antibodies as used in ExPLEX and followed the vendor-recommended protocol (R&D Systems) for ELISA. Capture antibody was diluted to a recommended concentration in PBS and added to the Maxisorp 96-well plate (Nunc) for overnight incubation at 4°C. After being washed three times with PBS-T, 2% BSA in PBS blocking solution was added to the plate for 2 h incubation at room temperature. Subsequently, serially diluted standard or samples were added to each well for at least 2 h incubation at room temperature. After the samples were discarded and washed three times with PBS-T, biotinylated detection antibodies (recommended concentrations, diluted in 0.1% BSA solution) were added to each well and incubated at room temperature for 2 h. Unbound antibodies were washed with PBS-T three times. High sensitivity streptavidin−HRP molecules (1:5000 diluted in 0.1% BSA solution) were added to the each well for 30 min at room temperature. After being washed out with PBS, a TMB solution was added to each well and incubated for 20 min, an equal volume of stopping solution (2 M H2SO4) was added, and the optical density was read at 450 nm.

#### qPCR

cDNA was synthesized from each bacterial RNA sample using the cDNA synthesis kit (AccuPower Rochetscript Cycle RT PreMix, Bioneer) under thermal cycling conditions (15°C for 20 sec, 50°C for 4 min, 60°C for 20 sec) for 10 cycles, followed by 95°C for 5 min. For quantitative real-time PCR, the cDNA derived from bacteria samples were added with the reverse transcription system (AccuPower^®^ Master Mix, Bioneer) and specific primers (**Supplementary Table 6**). The reaction mixtures were then applied to the Exicycler™ 96 Real-Time Quantitative Thermal Block and thermal cycling was carried out for 40 cycles at conditions of 94°C for 5 sec, annealing at 58°C for 25 sec, and extension at 72°C for 30 sec. Ct values were obtained using the Exicycler3 software (provided by the manufacturer), and relative amounts of target nucleic acid were calculated based on the Ct values.

### 13. Clinical samples

#### Plasma samples

Plasma samples were obtained from patients with H3K27M-DMG treated with ONC201. For patients treated at the University of Michigan, written informed consent and assent were obtained under an Institutional Review Board–approved protocol for the Sample Repository for Pediatric Hematology/Oncology (HUM00123426). Peripheral blood (6–8 mL) was collected in vacutainer tubes at time points coinciding with routine clinical blood draws. Samples were processed by centrifugation to isolate plasma, which was aliquoted (0.4 mL) and stored at −80 °C until EV analysis. Healthy, disease-free plasma controls from single donors were commercially acquired from Innovative Research and STEMCELL Technologies and stored at −80 °C until use. All plasma samples were de-identified prior to downstream processing. Enrollment numbers and patient information for the cohort used in this study are summarized in **Supplementary Table 1**.

#### MRI/RAPNO and tumor measurements

Tumor measurements for patients enrolled in the phase 1 ONC201 clinical trial (NCT03416530) were provided by study investigators and assessed using modified RANO response criteria. For contrast-enhancing disease, bi-dimensional measurements were performed on MRI scans using lesions with clearly defined margins, a minimal diameter of ≥ 1 cm, and visibility on at least two axial slices separated by ≥ 5 mm without skip. When contrast-enhancing disease was not measurable, bi-dimensional measurements were obtained from T2/FLAIR images. Tumor cross-sectional area was calculated from the axial slice exhibiting the maximal tumor diameter as the product of this diameter and the perpendicular diameter in the same plane. For correlation with EV mRNA measurements, MRI scans were selected from the imaging study closest in time to plasma sample collection.

### 14. Statistics & Reproducibility

Statistical analyses and data plotting were performed in GraphPad Prism 10. For correlations, the linear least squares fitting was performed at the 95% confidence level, and the Pearson correlation coefficient (*r*) was used to quantify the correlations between different variables.

Group differences were tested using the *t*-test for two groups and analysis of variance (ANOVA) with post hoc analysis for more than two groups. All tests were two-sided, and *p* < 0.05 was considered statistically significant.

## DATA AVAILABILITY

Source data underlying all figures are provided with the paper.

## ACKNOWLEDGEMENTS

We acknowledge the following grant support: V Scholar Award (V2023-002) funded by the V Foundation to J.M.; NIH R35GM157070 and NSF 2441718 supporting J.M.’s effort in part.

## AUTHOR CONTRIBUTION

J.M. and C.J.K. conceived the study. C.K. and J.K. performed the experiments. J.C. developed the DL algorithms. K.B. provided supervision of J.C. B.L., A.P., and C.J.K. coordinated the collection and processing of clinical samples and associated data. C.K., J.K., C.J.K., and J.M. analyzed the data. C.K., J.K., and J.M. wrote the manuscript, and all authors reviewed and edited the final version.

## COMPETING INTERESTS

The authors declare that they have no competing interests.

## Notes

### Competing Interest Statement

The authors have declared no competing interest.

