## Supplementary materials for "Tumor extracellular vesicle RNA profiling predicts treatment response in pediatric diffuse midline glioma"

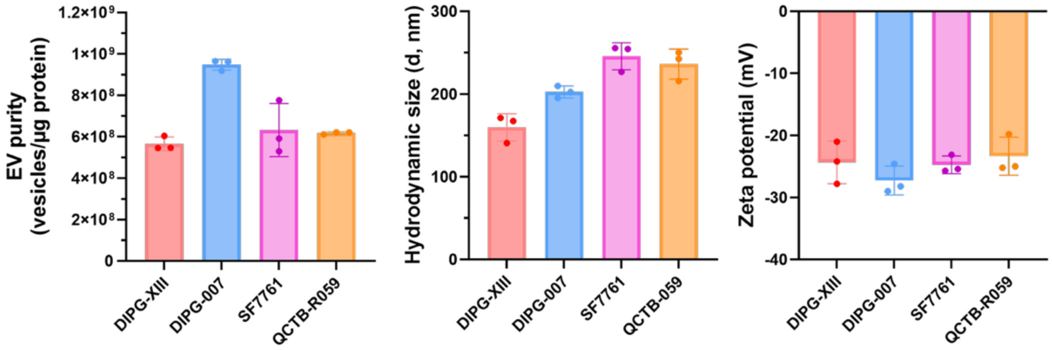


**Supplementary Fig. 1. Physical characterization of isolated DMG-EV.** EV purity, hydrodynamic size and zeta potential of DMG EV were characterized.

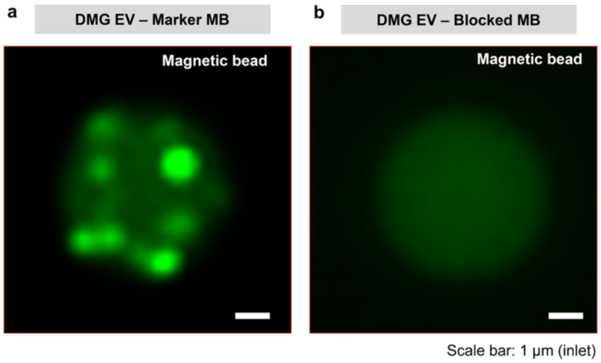


**Supplementary Fig. 2. Selective isolation of DMG-EVs through DMG-pan MB.** (a) BT245-derived EVs labeled with PKH67 being captured through immuno-interaction with DMG-pan MB. (b) In contrast, the IgG control MB, which was introduced with an IgG isotype control, did not capture the EVs.

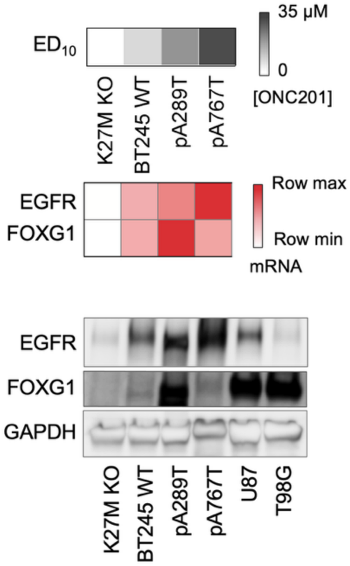


**Supplementary Fig. 3**. **EGFR and FOXG1 expression and ONC201 sensitivity in BT245-derived H3K27M-mutant DMG models.** EGFR and FOXG1 levels were assessed by qPCR and western blotting in genetically engineered BT245-derived models with distinct in vitro ONC201 sensitivities.

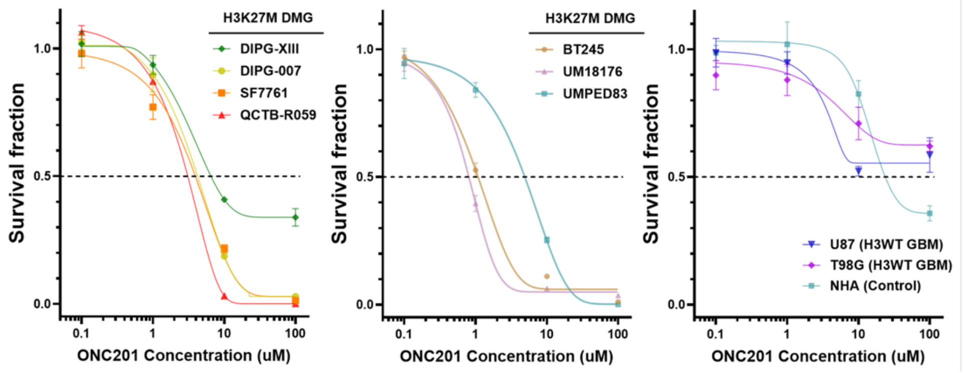


| Cell line | $IC_{50}$ (μM) | $E_{10}$ |
|---|---|---|
| UM18176 | 0.8 | 6.4 |
| BT245 | 1.1 | 11.1 |
| QCTB-R059 | 2.3 | 3.2 |
| SF7761 | 4 | 21.6 |
| DIPG-007 | 4.1 | 18.7 |
| UMPED83 | 5 | 25.4 |
| DIPG-XIII | 6.5 | 33.9 |
| U87 | - | 58.6 |
| T98G | - | 71 |
| NHA | 25.0 | 78 |

**Supplementary Fig. 4**. **Dose response of DMG cells when treated with ONC201.** Human DMG cells were treated with varying doses of ONC201, and their viability was determined. Cell lines were classified into sensitive (DIPG7, SF7761, QCTB-R059, BT245, UMPED83, UM18176; ED10 < 30 μM) and resistant (all other cell lines) according to their respective drug response. ED10, 10% effective dose.

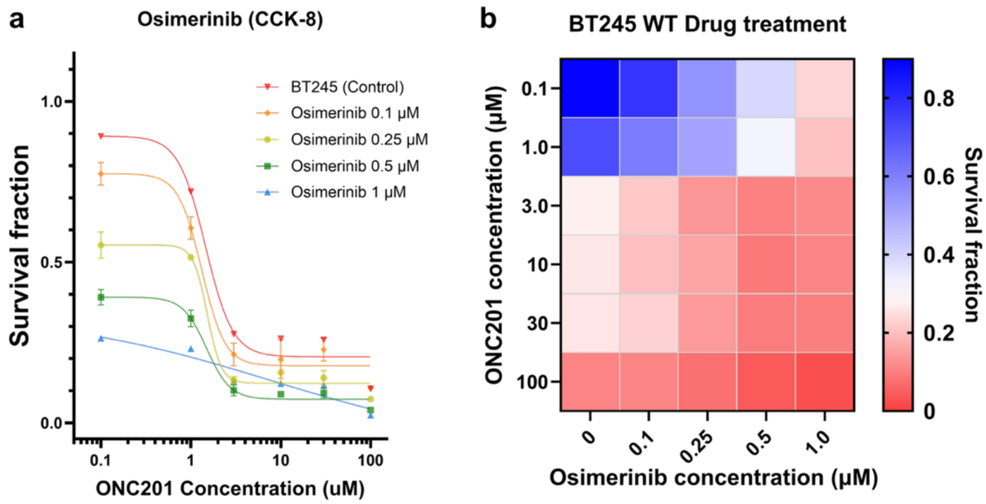


**Supplementary Fig. 5. Dose response of BT245 when treated with ONC201 and Osimerinib.** (a) BT245 was treated with varying doses of ONC201 and Osimerinib, which is EGFR inhibitor (EGFRi). (b) Heatmap image of viability via cocktail treatment. The greater the inhibition of EGFR by osimertinib, the greater the response to ONC201.

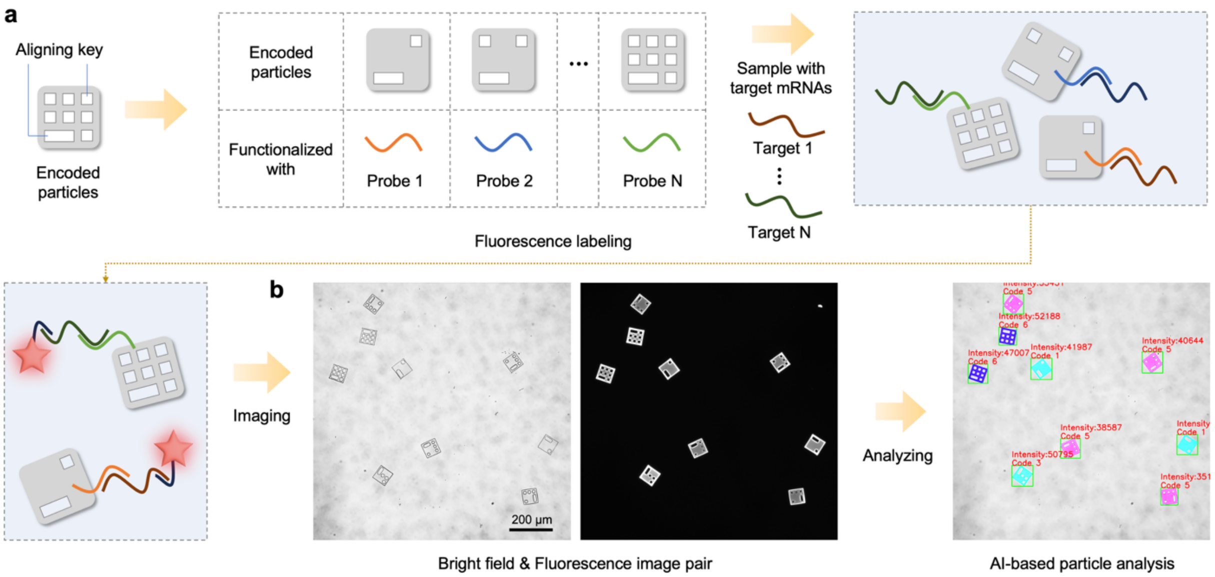


**Supplementary Fig. 6. Schematic overview of ExPLEX assay workflow. (a)** Shape-encoded hydrogel particles are functionalized with specific single-stranded DNA capture probes. A pooled mixture of encoded particles is incubated with the sample to enable simultaneous capture of multiple target mRNAs. Following fluorescence labeling, target-bound particles are prepared for imaging. **(b)** Representative field of view (FOV) showing bright-field (BFM) and fluorescence (FLM) images acquired using a 10× objective, along with the corresponding AI-based analysis output. The analysis includes automated particle detection, code decoding, and fluorescence intensity quantification.

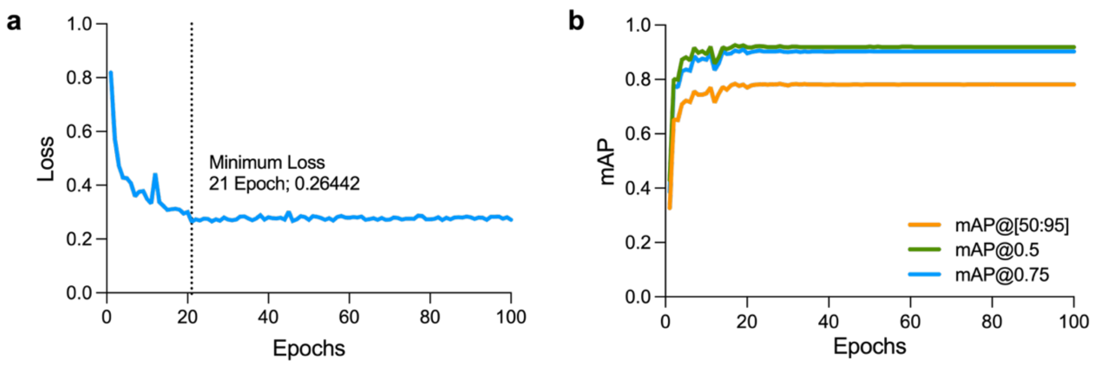


**Supplementary Fig. 7. Deep learning-based particle decoding for ExPLEX analysis.** (a) Validation loss versus training epoch for the Mask R-CNN model; the final model was selected at epoch 21, corresponding to the minimum loss. (b) Mean average precision (mAP) as a function of training epoch evaluated at different intersection-over-union (IoU) thresholds, demonstrating robust performance across stringent detection criteria.

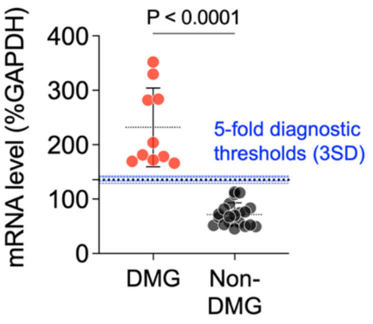


**Supplementary Fig. 8. Robustness of H3K27M positivity threshold assessed by 5-fold cross-validation.** The black dotted line indicates the overall threshold defined as mean + 3 × SD of the non-DMG control group, and the blue dashed lines show the thresholds obtained from each of the five folds.

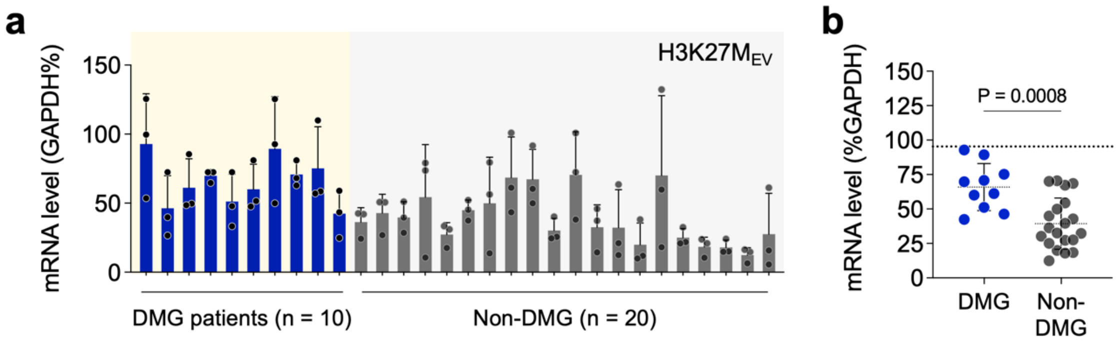


**Supplementary Fig. 9. EV analyses via conventional qPCR for differentiation of DMG patients from non-DMG control groups. (a)** Measurement of EV H3K27M mRNA levels measured by qPCR in plasma samples. **(b)** EV H3K27M mRNA levels were higher in DMG patients than in non-DMG controls (two-sided t-test, P = 0.0008). The dotted line indicates the preliminary positivity threshold, defined as mean + 3 × SD of the non-DMG control group.

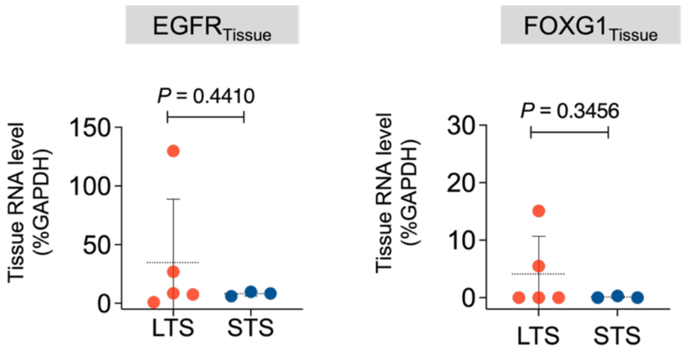


**Supplementary Fig. 10. Tissue analysis for ONC treatment response.** Measurement of mRNA for EGFR and FOXG1 as predictive markers for ONC-201 treatment response in tissue samples. (P = 0.4410 for EGFR and P = 0.3456 for FOXG1, tow-sided t-test).

**Supplementary Table 1**. Patient information. (STS, short-term survivor; LTS, long-term survivor)

| PATIENT ID | SAMPLE TYPE | COLLECTION TIME POINT | | AGE AT DX- (DAYS) | SEX | PRIMARY LOCATION | ONC THERAPY | ONC START | ONC STOP | PFS | SURVIVAL CLASS |
|---|---|---|---|---|---|---|---|---|---|---|---|
| | | Pre-TX (Days) | Post-TX (Days) | | | | | | | | |
| UMPED175 | Plasma | 0 | 191 | 2801 | M | Pons | ONC206 | 2823 | 3215 | 303 | STS |
| UMPED213 | Plasma | 0 | 112 | 1211 | M | Pons | ONC201/206 | 1317 | 1535 | 316 | LTS |
| UMPED193 | Plasma | 0 | 141 | 6919 | M | Pons | ONC201 | 7070 | 7575 | 375 | LTS |
| UMPED217 | Plasma | 0 | 175 | 6604 | M | Pons | ONC201 | 6714 | 6910 | 305 | STS |
| UMPED209 | CSF | 0 | 145 | 4991 | F | Pons | ONC201 | 5105 | 5250 | 223 | STS |
| UMPED54 | Plasma | 0 | 1722 | 2365 | F | Thalamus | ONC201 | 2451 | 4201 | 1805 | LTS |
| UMPED207 | Plasma | 0 | 147 | 4417 | M | Pons | ONC201 | 4500 | 4658 | 253 | STS |
| UMPED104 | Plasma | 0 | 1609 | 3228 | F | Thalamus | ONC201 | 3334 | Current | 2011 | LTS |
| UMPED156 | Plasma | 0 | 35 | 2168 | F | Pons | ONC201 | 2267 | 2337 | 141 | STS |
| UMPED192 | Plasma | 0 | 252 | 7447 | M | Thalamus | ONC201 | 7504 | 7886 | 392 | LTS |

**Supplementary Table 2.** Cell lines information.

| CELL LINE | NOTES | SOURCE | CELL TYPE | MEDIA |
|---|---|---|---|---|
| DIPG-7 | K27M-mutant DMG | UMHS | Suspension | TSM |
| DIPG-13 | K27M-mutant DMG | UMHS | Suspension | TSM |
| SF7761 | K27M-mutant DMG | UMHS | Suspension | TSM |
| QCTB-R059 | K27M-mutant DMG | UMHS | Suspension | TSM |
| BT245 | K27M-mutant DMG | UMHS | Suspension | TSM |
| UMPED83 | K27M-mutant DMG | UMHS | Suspension | TSM |
| UM18671 | K27M-mutant DMG | UMHS | Suspension | TSM |
| U87 | H3WT glioblastoma | ATCC | Adherent | DMEM media |
| T98G | H3WT glioblastoma | ATCC | Adherent | DMEM media |
| NHA | Primary human astrocyte | ScienCell | Adherent | DMEM media |

**Supplementary Table 3.** Cell culture media.

| MEDIA | REAGENT | VENDOR | WORKING CONCENTRATION |
|---|---|---|---|
| Tumor stem media (TSM) | DMEM/F12 | Gibco | - |
| | B-27 Supplement without vitamin A (50X) | Gibco | 1X |
| | Human epidermal growth factor (H-EGF) | Peprotech | 20 ng/mL |
| | Human Fibroblast growth factor (H-FGF) | Peprotech | 20 ng/mL |
| | Human platelet-derived growth factor (PDGF)-AA | Peprotech | 10 ng/mL |
| | Human platelet-derived growth factor (PDGF)-BB | Peprotech | 10 ng/mL |
| | Antibiotic-antimycotic (100X) | Gibco | 1X |
| | Normocin | InvivoGen | 1 mg/mL |
| | Heparin | StemCell Technologies Inc. | 2 μg/mL |
| DMEM media | DMEM | Gibco | - |
| | Fetal bovine serum (FBS) | Corning | 10% |
| | Penicillin-Streptomycin | Gibco | 50 U/mL |
| Mycoplasma detection kit | MycoAlert Mycoplasma Detection Kit | Lonza | - |
| | MycoStrip - Mycoplasma Detection Kit | InvivoGen | - |

**Supplementary Table 4.** Antibody information. WB = Western blot, MB = Magnetic bead.

| COMPONENT | TARGET | CLONE | VENDOR | NOTES |
|---|---|---|---|---|
| EV marker | CD9 | H19a | Biolegend | 1:500 for WB, MB |
| | CD63 | Ts63 | Thermofisher | 1:500 for WB, MB |
| | CD81 | 5A6 | Biolegend | 1:500 for WB, MB |
| | TSG101 | W19019A | Biolegend | 1:1000 for WB |
| | Arf6 | D12G6 | CST | 1:1000 for WB |
| Cancer | EGFR | A19002A | CST | 1:1000 for WB; MB |
| | HER2 | 29D8 | CST | 1:1000 for WB |
| | GD2 | 14G2a | CST | 1:1000 for WB; MB |
| | PD-L1 | 29E.2A3 | CST | 1:1000 for WB |
| | IL-13Rα2 | E7U7B | CST | 1:1000 for WB |
| | | A21071B | Biolegend | MB |
| Others | ApoA | 5F4 | CST | 1:1000 for WB |
| | ApoB | EPR2914 | Abcam | 1:1000 for WB |
| | Calnexin | C5C9 | CST | 1:1000 for WB |
| | Albumin | N/A | CST | 1:1000 for WB |
| | GAPDH | 14C10 | CST | 1:1000 for WB |
| | β-actin | 2F1-1 | Biolegend | 1:1000 for WB |
| Secondary antibodies | Anti-rabbit IgG antibody, HRP | N/A | CST | 1:2000-1:5000 for WB |
| | Anti-mouse IgG antibody, HRP | N/A | CST | 1:2000-1:5000 for WB |
| IgG control | Mouse IgG isotype control | MG1-45 | Biolegend | MB |
| | Rabbit IgG isotype control | Poly29108 | Biolegend | MB |

**Supplementary Table 5.** Sequences for probe targeting EV RNAs for ExPLEX assay.

| TARGET | CAPTURE PROBE (5'->3') | DETECTION PROBE (5'->3') |
|---|---|---|
| GAPDH | CGCTGTTGAAGTCAGAGGA-A6-Thiol | Biotin-A6-CCACCCTGTTGCTGTAGCCAA |
| H3F3A mutation | TCATGCGAGCGGCTTTTG-A6-Thiol | Biotin-A6-TTCACGGAGCGCCACAGTA |
| EGFR | GCCTGTCGTCCGGTCTGG-A6-Thiol | Biotin-A6-TCCGGCTCTCCCGATCAAT |
| FOXG1 | GGAAATCTGGCGGCTCTT-A6-Thiol | Biotin-A6-TGTTCTCAAGGTCTGCGTC |

**Supplementary Table 6.** Primer sequences for PCR.

| TARGET | FORWARD PRIMER (5'->3') | REVERSE PRIMER (5'->3') |
|---|---|---|
| GAPDH | GTCTCCTCTGACTTCAACAGC G | ACCACCCTGTTGCTGTAGCCA A |
| H3F3A mutation | CAAAAGCCGCTCGCAT | TTCACGGAGCGCCACAGTA |
| EGFR | CCAGACCGGACGACAGG | TCCGGCTCTCCCGATCAATA |
| FOXG1 | AAGAGCCGCCAGATTTCCAT | TGTTCTCAAGGTCTGCGTCC |
| TP53 | CCTCAGCATCTTATCCGAGTGG | TGGATGGTGGTACAGTCAGAG C |
| ATF4 | TCCGAATGGCTGGCTGTGG | AGTGTAGTCTGGCTTCCTATCTCC |
| CD63 | AACCACACTGCTTCGATCCT | AATCCCACAGCCCACAGTAA |
| DAPK2 | TCCTGGATGGGGTGAACTAC | CAGCTTGATGTGTGGAATGG |
| CPLX2 | GGAGAGAGGCCAAGATATTAAG | CGTACCAGCAGACAGATTATT |
| CKMT2 | GGAGAGAGGCCAAGATATTAAG | CGTACCAGCAGACAGATTATT |
| RPS6KB2 | CTTCCAGACTGGTGGCAAACTCTA | CAGCGTGATCTCAGCCAGGTA |
